# Education and coronary heart disease: a Mendelian randomization study

**DOI:** 10.1101/106237

**Authors:** Taavi Tillmann, Julien Vaucher, Aysu Okbay, Hynek Pikhart, Anne Peasey, Ruzena Kubinova, Andrzej Pajak, Abdonas Tamosiunas, Sofia Malyutina, Krista Fischer, Giovanni Veronesi, Tom Palmer, Jack Bowden, George Davey Smith, Martin Bobak, Michael V Holmes

**Affiliations:** Department of Epidemiology & Public Health, University College London, UK; Department of Internal Medicine, Lausanne University Hospital, Switzerland; Department of Applied Economics, Erasmus University, Netherlands; Department of Environmental Health Monitoring System, National Institute of Public Health, Prague, Czech Republic; Department of Epidemiology and Population Studies, Institute of Public Health, Jagiellonian University, Krakow, Poland; Institute of Cardiology, Lithuanian University of Health Sciences, Kaunas, Lithuania; Institute of Internal and Preventive Medicine, Novosibirsk, Russia; Novosibirsk State Medical University, Novosibirsk, Russia; Estonian Genome Center, University of Tartu, Estonia; Research Center in Epidemiology and Preventive Medicine, University of Insubria, Varese, Italy; Department of Mathematics and Statistics, Lancaster University, Lancaster, UK; Medical Research Council Integrative Epidemiology Unit at the University of Bristol, UK; Clinical Trial Service Unit & Epidemiological Studies Unit, Nuffield Department of Population Health, University of Oxford, UK; Medical Research Council Population Health Research Unit at the University of Oxford, UK

**Author notes:** Correspondence to Department of Epidemiology & Public Health, University College London, 1-19 Torrington Place, WC1E 7HB, UK. And Julien Vaucher, Service of Internal Medicine, Lausanne University Hospital, Bugnon 46, 1011 Lausanne, Switzerland. =contributed equally (Taavi Tillmann, Julien Vaucher).

**Keywords:** Socioeconomic Factors, Educational status, Cardiovascular disease, Myocardial ischemia, Coronary artery disease, Myocardial infarction, Prospective studies, Mendelian

## Abstract

**Objectives:** To determine whether educational attainment is a causal risk factor in the development of coronary heart disease.

**Design:** Mendelian randomization study, where genetic data are used as proxies for education, in order to minimize confounding. A two-sample design was applied, where summary level genetic data was analysed from two publically available consortia.

**Setting:** In the main analysis, we analysed genetic data from two large consortia (CARDIoGRAM and SSGAC), comprising of 112 cohorts from predominantly high-income countries. In addition, we also analysed genetic data from 7 additional large consortia, in order to identify putative causal mediators.

**Participants:** The main analysis was of 589 377 men and women, predominantly of European origin.

**Exposure:** A one standard deviation increase in the genetic predisposition towards higher education (i.e. 3.6 years of additional schooling). This was measured by 162 genetic variants that have been previously associated with education.

**Main outcome:** Combined fatal and nonfatal coronary heart disease (63 746 events).

**Results:** 3.6 years of additional education lowered the risk of coronary heart disease by a third (odds ratio = 0.67, 95% confidence interval [CI], 0.59 to 0.77, p=0.01). Equivalent increases in education were also causally associated with reductions in smoking, BMI and improvements in blood lipid profiles.

**Conclusions:** More time spent in education is causally associated with a large reduction in the risk of coronary heart disease. This may be partly explained by changes to smoking, BMI and a blood lipids. These findings offer support for policy interventions that increase education, in order to also reduce the burden of cardiovascular disease.

## What this paper adds

### What is already known on this subject

Numerous observational studies have found that people with longer educational attainment develop less coronary heart disease. However it is not clear whether this association is causal, partly since randomised controlled trials are not feasible in this area. No prior studies have applied the Mendelian randomization method to investigate how exposure to socioeconomic risk factors might causally change the risk of disease occurrence.

### What this study adds

Our study suggests that if young people were to increase the number of years they spend in the educational system, then this is likely to substantially lower their risk of subsequently developing coronary heart disease. Our study should stimulate further policy discussion about increasing educational attainment in the general population, in order to improve population health.

## INTRODUCTION

Coronary heart disease (CHD) is the leading cause of death globally. While the causal effects of risk factors like smoking, blood pressure and LDL-cholesterol are generally accepted and reflected in disease prevention strategies, substantial uncertainty still surrounds other potential factors. Decades of observational studies have consistently associated socioeconomic factors such as higher education with decreased risk of CHD.^1–4^ However, given the implausibility and consequent absence of randomized controlled trials in this field, it is possible that this association does not stem from an underlying causal effect, but arises due to the methodological flaws of observational research.^5 6^ Clarifying whether the association between education and CHD is causal or not has widespread implications for our understanding of the aetiology of CHD, and the development of novel population-based approaches to its prevention.

Mendelian randomization analysis relies on genetic variants that are associated with a risk factor (e.g. education), to make causal inferences about how environmental changes to the same risk factor would alter the risk of disease (e.g. CHD).^7^ Genetic variants associated with a risk factor are independent from confounders that may otherwise cause bias, as genetic variants are randomly allocated before birth.^8^ This, as well as the non-modifiable nature of the genetic variants, provides an analogy to trials, where exposure is allocated by random and is non-modifiable by disease.^8^ By comparing the risk of disease across the genotype groups, this allows causal effects to be determined with substantially less bias than that from observational epidemiology. Recent methodological developments, including Egger Mendelian randomization (MR-Egger), weighted median MR and multivariable MR, can be employed as sensitivity analyses to additionally investigate any pleiotropic effects of the genetic variants (i.e., when genetic variants associate with one or more pathways than with the risk factor under analysis, which can distort the causal effect estimate) **(Figure S1).**^9–11^

Here, we first updated observational estimates of the association between education and incidence of coronary heart disease from several large studies **(Table S1)**. A recent genome-wide association study (GWAS) from the Social Science Genetic Association Consortium identified a large number of independent genetic variants (i.e., single-nucleotide polymorphisms [SNPs]) associated with educational attainment.^12^ Thus, we used two sets of SNPs associated with education attainment and retrieved the same CHD-associated SNPs from the CARDIoGRAMplusC4D Consortium to perform Mendelian randomization analyses, investigating whether individuals with a genetic predisposition towards higher education have a lower risk of CHD.^13^ We checked the robustness of our findings across a range of sensitivity analyses. We finally investigated whether genetic liability for CHD is associated with educational outcomes.

## METHODS

### Observational association between education and CHD

Throughout all analyses, CHD was defined as a composite of myocardial infarction, acute coronary syndrome, chronic stable angina or coronary stenosis of >50%, or coronary death. In observational analysis, we used a combination of cross-sectional and prospective data, collected between 1983-2014 **(Table S1).** For prevalent CHD cases, we analysed 43,611 participants (1,933 cases) from the National Health and Nutrition Examination Surveys (NHANES) **(Tables 1** and **S1, Figure S2)** (http://www.cdc.gov/nchs/nhanes).^14^ For incident cases, we analysed 23,511 participants (632 CHD cases) from the Health, Alcohol and Psychosocial factors In Eastern Europe (HAPIEE) study and combined this with published estimates from 97,048 participants (6,522 cases) of the MOnica Risk, Genetics, Archiving and Monograph (MORGAM) study in Europe (see **Table S1** for case definitions, and **Figures S3-S4**).^16^

#### Genetic variants associated with education

SNPs associated with educational attainment were retrieved from a recent genome-wide association study, involving 405,072 individuals of European ancestry **(Table 1)**.^12^ For our main analysis, we used 162 independent SNPs (linkage disequilibrium, *r*^2^<0.1) associated with education across the discovery and replication cohorts at a combined GWAS significance threshold (P<5.10^−8^) **(Supplementary Methods, Table S2-S3** and **Figure S5-S6).** For our secondary analysis, we used another set of 72 independent SNPs (linkage disequilibrium, *r*^2^<0.1) that were associated with education in the discovery cohort (293,723 participants; P<5.10^−8^) and directionally consistent in the replication cohort **(Supplementary Methods, Table S3-S4,** and **Figure S5** and **S7).**

**Table 1.**
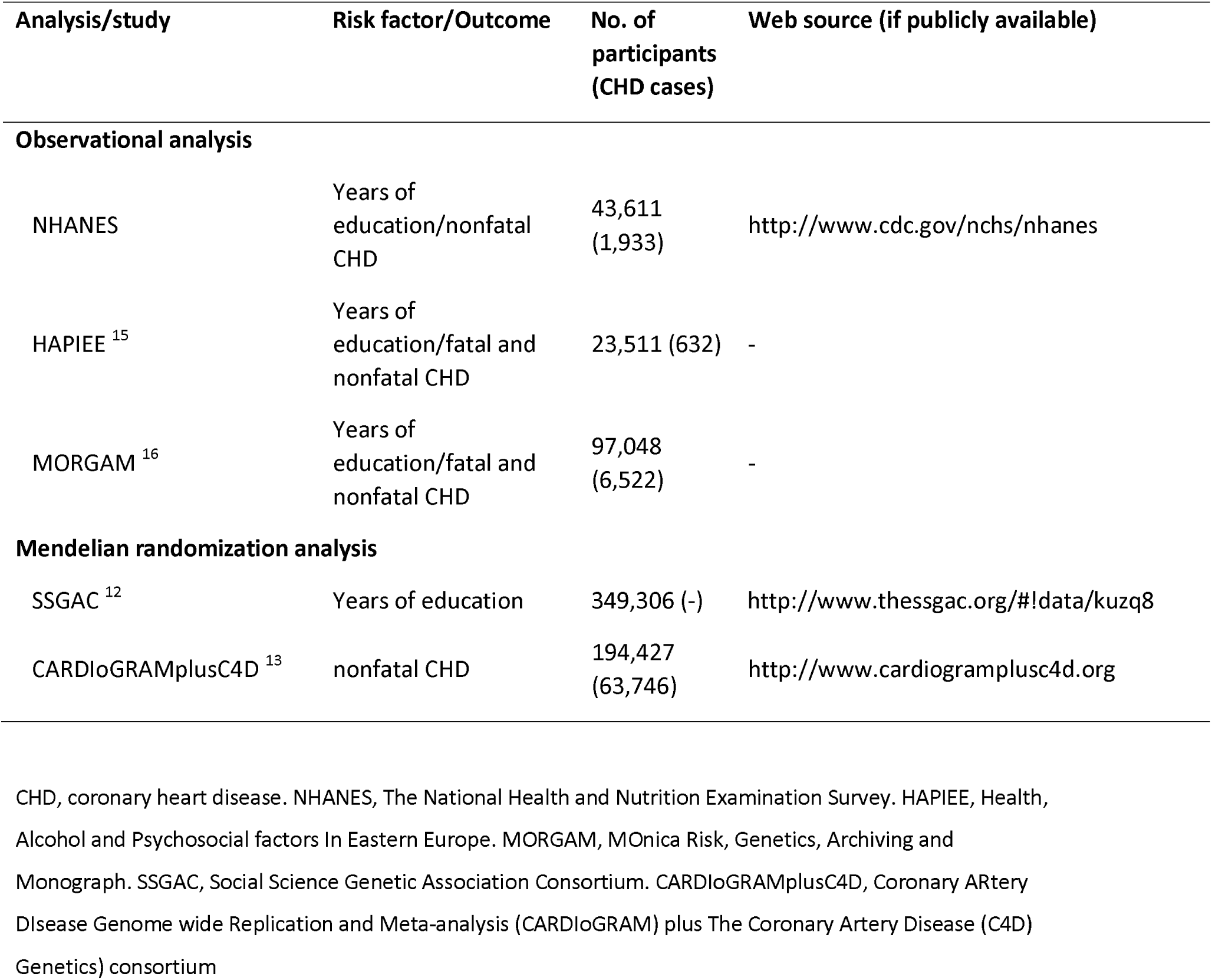
Details of the studies and datasets included in the analysis.

| Analysis/study | Risk factor/Outcome | No. of participants (CHD cases) | Web source (if publicly available) |
| --- | --- | --- | --- |
| **Observational analysis** |  |  |  |
| NHANES | Years of education/nonfatal CHD | 43,611 (1,933) | http://www.cdc.gov/nchs/nhanes |
| HAPIEE <sup>15</sup> | Years of education/fatal and nonfatal CHD | 23,511 (632) | - |
| MORGAM <sup>16</sup> | Years of education/fatal and nonfatal CHD | 97,048 (6,522) | - |
| **Mendelian randomization analysis** |  |  |  |
| SSGAC <sup>12</sup> | Years of education | 349,306 (-) | http://www.thessgac.org/#!data/kuzq8 |
| CARDIoGRAMplusC4D <sup>13</sup> | nonfatal CHD | 194,427 (63,746) | http://www.cardiogramplusc4d.org |

CHD, coronary heart disease. NHANES, The National Health and Nutrition Examination Survey. HAPIEE, Health, Alcohol and Psychosocial factors In Eastern Europe. MORGAM, MONica Risk, Genetics, Archiving and Monograph. SSGAC, Social Science Genetic Association Consortium. CARDIoGRAMplusC4D, Coronary ARtery Disease Genome wide Replication and Meta-analysis (CARDIoGRAM) plus The Coronary Artery Disease (C4D) Genetics) consortium

### Genetic variants associated with CHD

For each education-associated SNP, we retrieved summary-level data from the Coronary ARtery Disease Genome wide Replication and Meta-analysis plus The Coronary Artery Disease Genetics Consortium (CARDIoGRAMplusC4D) comprising 63,746 CHD cases and 130,681 controls **(Supplementary Methods, Table 1, and Figures S8-S9)**.^13^

### Statistical analyses

Cox proportional hazards and logistic regressions were used to calculate observational estimates for incident and prevalent cases, respectively. Conventional Mendelian randomization analyses were performed by regressing the SNP-education associations with the SNP-CHD associations (outcome), where each SNP was one data point **(Supplementary Methods)**.^9^

Sensitivity analyses (MR-Egger and weighted median MR) were used to investigate to what degree pleiotropic effects might bias the Mendelian randomization causal estimates.^9 10^ In addition, we also applied Mendelian randomization to investigate whether genetic predisposition to higher education could lead to improvements in the established cardiovascular risk factors **(Supplementary Methods** and **Table S5).** Once such associations were identified, we further adjusted for their causal effects using multivariate MR.^11^ Finally, to check for whether genetic risk for coronary events might be a causal factor for educational attainment, we performed Mendelian randomization in the opposite direction **(Supplementary Methods** and **Table S6).**

### Patient involvement

Patients were not involved in the design or implementation of this study. There are no specific plans to disseminate the research findings to participants, but findings will be returned back to the original consortia, so that they can consider disseminating further.

## RESULTS

### Observational analyses

Based on NHANES data, each additional 3.6 years of education (1-SD) was associated with 27% lower odds of CHD (odds ratio, 0.73; 95% confidence interval [CI], 0.68 to 0.78) **(Figure 2).** In prospective analyses, 3.6 years of additional education was associated with a 20% lower risk of CHD in the HAPIEE and MORGAM studies, with a pooled hazard ratio of 0.80 (95% CI, 0.76 to 0.83) **(Figure S10)**.^15 16^ These observational estimates were robust to sensitivity analyses accounting for different case definitions, different age at first event, and confounding by other measures of socioeconomic position **(Table S7).**

### Causal association of education on CHD

When examining SNPs across the entire genome **(Supplementary Methods),** there was strong evidence for a negative genetic correlation between education and CHD (rg=−0.324; p-value=2.1 x 10^−12^).^17^

Conventional Mendelian randomization analysis using 162 SNPs found that a 1-SD increase in education was associated with a 33% reduction in the risk of CHD (odds ratio, 0.67; 95% CI, 0.59 to 0.77) **(Figure 1** and **S11).** Sensitivity analyses using MR-Egger and weighted median Mendelian randomization yielded similar results in terms of direction and magnitude **(Figures 1** and **S12-S13** and **Table S8),** suggesting that our findings were not biased from presence of pleiotropic effects. The secondary analysis using a set of 72 SNPs yielded consistent results in terms of direction and magnitude **(Figures 1 and S14-S16, Table S8)**.^18^

**Figure 1.**
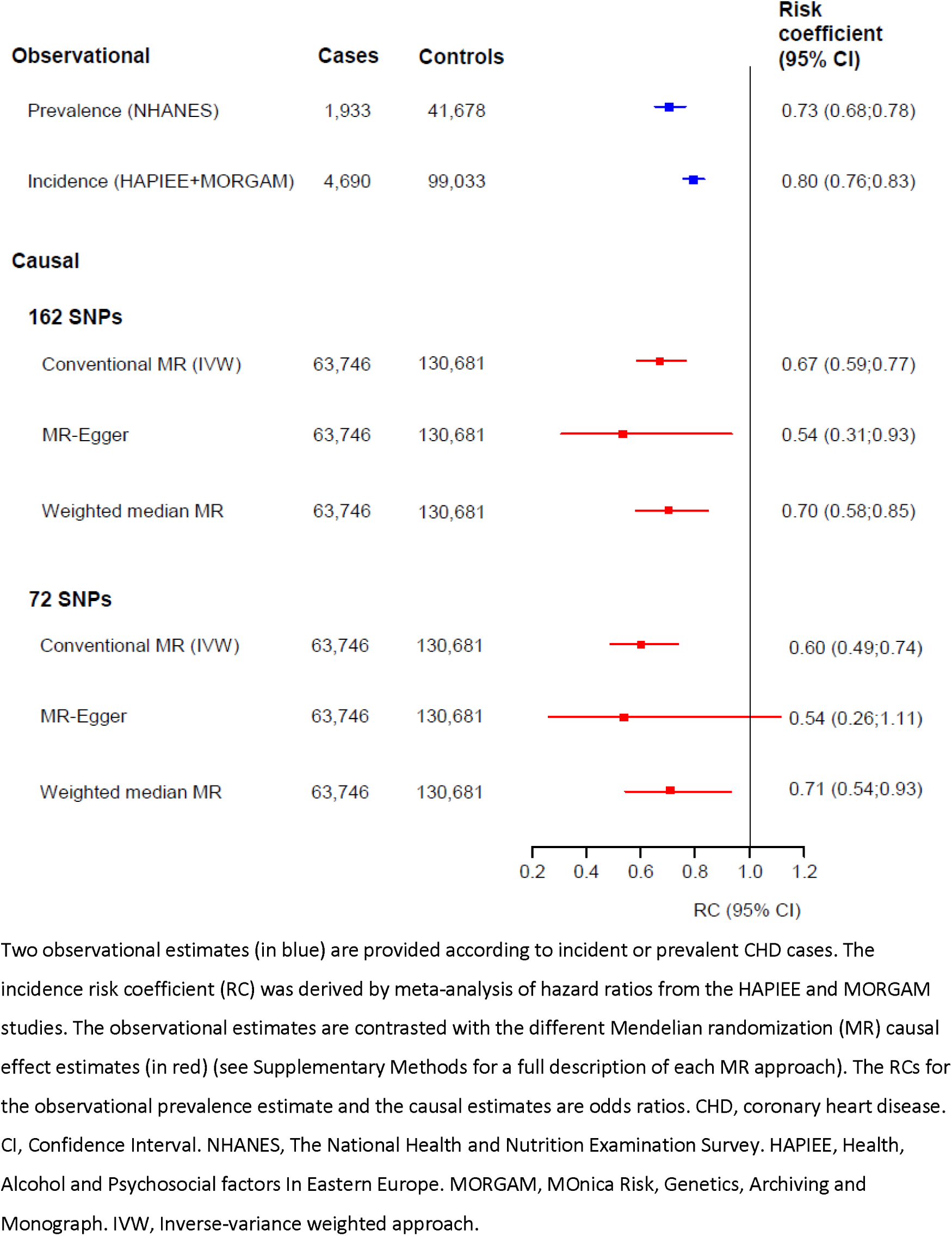
Comparison of observational (blue) and causal (red) estimates for risk of coronary
heart disease, per 3.6 years of educational attainment

### Causal association of education on cardiovascular risk factors

To identify potential risk factors that could mediate or confound the association between education and CHD, we investigated whether a genetic predisposition towards higher education was associated with established cardiovascular risk factors **(Table 2** and **S5).** A 1-SD increase in education was causally associated with a 35% reduction in odds of smoking, a 0.17 kg/m^2^ reduction in BMI, a 0.14 mmol/l reduction in triglycerides and a 0.15 mmol/l increase in HDL-cholesterol **(Table 2).** In exploratory analyses, inclusion of genetic associations with smoking, BMI, triglycerides and HDL-cholesterol did not change the primary association between education and CHD **(Tables 3** and **S9).**

**Table 2.** Causal effects of a 1-SD increase in education on selected cardiovascular risk factors

| Outcome | Estimate (95% CI) per<br>1-SD increase in<br>education (~3.6 years) | P-value |
| --- | --- | --- |
| Binary traits | (estimate is Odds Ratio) |  |
| **Smoking status** | **0.65 (0.54; 0.79)** | **0.001** |
| Diabetes mellitus, type 2 | 0.75 (0.56; 1.01) | 0.057 |
| Continuous traits (units) | (estimate is beta) |  |
| Systolic Blood Pressure (mm Hg) | -1.36 (-2.85; 0.12) | 0.075 |
| Diastolic Blood Pressure (mm Hg) | -0.23 (-1.22; 0.76) | 0.645 |
| LDL-cholesterol (mmol/L) | -0.03 (-0.10; 0.05) | 0.513 |
| **HDL-cholesterol (mmol/L)** | **0.15 (0.07; 0.23)** | **0.001** |
| **Triglycerides (mmol/L)** | **-0.14 (-0.22; -0.06)** | **0.001** |
| Glucose (mmol/L) | -0.02 (-0.08; 0.03) | 0.441 |
| **Body Mass Index (kg/m<sup>2</sup>)** | **-0.17 (-0.26; -0.08)** | **0.001** |
| Height (cm) | 0.06 (-0.03; 0.16) | 0.208 |

**Table 3.** Multivariable Mendelian randomization analyses showing the causal effect of 1-SD increase in education on the risk of CHD

| **Risk factor adjusted for** | **Odds Ratio of CHD**<br>**(95% CI) per 1-SD**<br>**increase in**<br>**education (~3.6**<br>**years)** | **P-value** | **Attenuation** |
| --- | --- | --- | --- |
| Nil (unadjusted) | 0.79 (0.66; 0.94) | 0.010 | reference |
| Tobacco (ever vs. never users) | 0.80 (0.66; 0.97) | 0.023 | 5% |
| Body Mass Index | 0.81 (0.67; 0.98) | 0.035 | 11% |
| Triglycerides | 0.76 (0.63; 0.92) | 0.006 | -16% |
| HDL-cholesterol | 0.81 (0.67; 0.98) | 0.033 | 11% |
| All 4 risk factors | 0.80 (0.65; 0.99) | 0.039 | 5% |

### Causal association of CHD on education

The reverse investigation, of whether genetic liabilities for risk of CHD are associated with educational outcomes, showed an absence of association **(Figures 2** and **S17-S19).**

**Figure 2.**
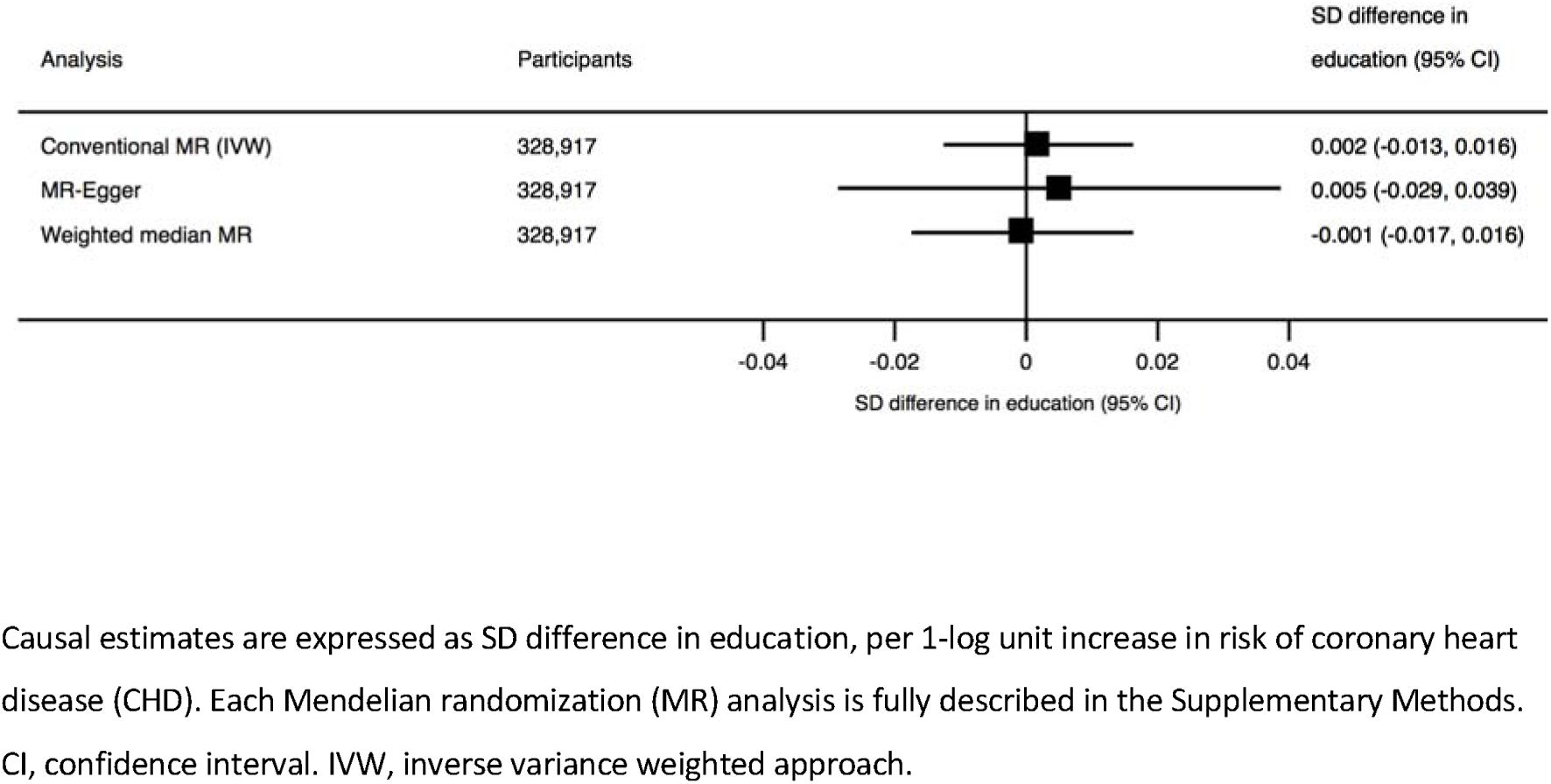
Association of genetic liabilities for risk of CHD and educational outcomes

## DISCUSSION

In this Mendelian randomization study, we found evidence that higher education is causally related to substantial reductions in the risk of CHD, consistent with observational estimates. More specifically, 3.6 years of increased education (similar to an undergraduate university degree) translates to about a one-third reduction in the risk of CHD.

Previous studies have attempted to clarify the causality of the relationship between education and CHD using approaches that overcome some of the limitations of observational epidemiology. These have primarily investigated the association between education and all-cause mortality. Few have investigated cardiovascular outcomes, and none have used a genetic approach using Mendelian randomization analysis. First, ecological instrumental variable approaches have compared mortality before and after changes to compulsory schooling laws. For example, by looking at mortality rates in countries before and after the introduction of national legislation that increased minimum education. In the UK, this increase in minimum education by one year was not associated with a change in mortality, but a similar increase in the Netherlands was associated with reduced mortality.^5 19^ Analyses of inter-state differences within USA initially suggested a large effect on mortality, but this effect disappeared when state-specific baseline trends were adjusted for.^20 21^ In Sweden, an intervention to extend compulsory schooling throughout a 13-year transition period was not associated with lower mortality for all age groups combined, but was associated with lower mortality in the younger age groups.^22^ Second, studies on twins have found differences between siblings in terms of development of chronic diseases when accounting for education, suggesting that confounding from environmental factors is unlikely to explain the observational associations.^23^-^25^ Third, studies including individual-level data from millions of siblings observed an association between education and mortality.^6 26^ Fourth, a recent study has reported a link between parental longevity and genetic markers for education in their offspring.^27^ However, causation and its direction were not tested.

The mechanisms that might mediate the association between education and CHD remain relatively unknown. Observational associations have found that the primary association attenuates by around 30-45% after statistical adjustment for health behaviours and conventional cardiovascular risk factors (including smoking, blood pressure and cholesterol). This suggests that these factors could account for around half of the association between education and CHD.^2 28^ In this study, we observed that education is causally related to smoking, BMI and blood lipids. Clarifying how much they mediate the causal association between education and CHD requires further investigation, e.g. by applying two-step Mendelian randomization to individual-level data. ^**29 30**^ Nonetheless, accounting for these factors in our multivariable Mendelian randomization analysis did not influence the primary association between education and risk of CHD, suggesting that conventional risk factors that we assessed may not completely account for the mechanism. Additional hypotheses for investigation could include education leading to improved use of healthcare services (from better health knowledge or fewer financial barriers in accessing care), better job prospects, income, material conditions, social ranking and/or diet - all factors associated education and CHD.^4^

Our study has important strengths. We investigated the causality of the association between an easily measured socioeconomic factor (i.e. education) and a common disease (i.e. coronary heart disease), using a genetic design to substantially reduce bias. Summary-level data from over half a million individuals gave our study the precision required to derive robust causal effect estimates, and to perform multiple sensitivity analyses. The consistency of findings across two sets of genetic instruments strengthens the confidence in our findings. We used recent state-of-the-art methodological developments to thoroughly explore pleiotropy our genetic variants, for which we found no evidence. However, our study also has some limitations. First, it is possible that the genetic variants associated with education may instead mark more generic biological pathways (such as vascular supply or mitochondrial function), which could simultaneously lead to increased educational attainment and reduced risk of CHD.^25 31^ In this scenario, interventions in education may not translate into lower disease occurrence. However, such a scenario is less likely to lead to the consistent set of results we found across all the sensitivity analyses. Second, it is possible that some of the genetic markers used were invalid. Nonetheless, even if all SNPs included in the analysis have pleiotropic effects, MR-Egger analyses should still give valid causal estimates, as long as the pleiotropic effects do not correlate with the effects on education. Third, we assumed the absence of dynastic effects, an assumption that is broken when parental genes associate with parental behaviours that directly cause a health outcome in the child. For example, parents with a genetic predisposition towards higher education may choose to feed their children a better diet. Fourth, our observational and genetic data originate predominantly from European origin samples in high-income countries. We are thus unable to generalize these estimates to other populations, particularly to low-income countries where cardiovascular diseases are less common. Fifth, we do not know whether increasing education for those of least education will be as cardioprotective as increasing education for those with above-average education. Nonetheless, the linear relationship in the observational data supports interventions framed within Rose’s prevention paradox: that more disease could be prevented by slight increases in education across its entire distribution, as opposed to targeting interventions only at high-risk subgroups.^33^ Sixth, we assumed that genetic predisposition towards a higher educational attainment causes the same behavioural and physiological consequences as environmentally-acquired changes to educational attainment. Although our study did not allow us to directly test the non-genetic component of education, these genetic findings are generally consistent with the various studies observing differences in mortality following policy change to schooling laws (as described above), as well as with observational estimates that measure the combined genetic and acquired components of education. Moreover, genetic predispositions to e.g. LDL-cholesterol and systolic blood pressure levels have recapitulated the direction of causality seen in environmentally-acquired changes to these risk factors from randomized controlled trials of pharmacological therapies, supporting the overall validity of the MR approach.^9 34^

In conclusion, this study suggests that a causal relationship exists between more time spent in education and a reduced risk of CHD. This relationship is only partly explained by traditional cardiovascular risk factors. Our findings add to the growing evidence base that increasing education is likely to lead to not only important societal benefits, but also substantial health benefits.

## Competing interests

All authors have completed the ICMJE uniform disclosure form at www.icmje.org/coi_disclosure.pdf and declare: no financial relationships with any organisations that might have an interest in the submitted work in the previous three years; no other relationships or activities that could appear to have influenced the submitted work.

## Contributors

Idea: TT. Genetic Data: AO. Observational Data: HP, AP, RK, AP, AT, SM, KF. Methods: MVH, GDS, JB, TP, GV. Analysis: JV, TT, MVH. Interpreted data: all co-authors. Wrote first draft: TT, JV, MVH. Revised manuscript for critical revisions: all co-authors. TT, JV and MVH are guarantors. TT, JV and MVH affirm that this manuscript is an honest, accurate, and transparent account of the study being reported; that no important aspects of the study have been omitted; and that any discrepancies from the study as planned have been explained.

## Acknowledgements

TT is funded by a Wellcome Trust Fellowship [106554/Z/14/Z]; JV is supported by the Swiss National Science Foundation (P2LAP3_155086); The HAPIEE study is supported by the Wellcome Trust [064947/Z/01/Z, WT081081]; the US National Institute on Aging [1R01 AG23522]; the MacArthur Foundation [Health and Social Upheaval network]; and the Russian Science Foundation [14-45-00030]; GDS works in the Medical Research Council Integrative Epidemiology Unit at the University of Bristol [MC_UU_12013/1]. The funders, had no role in the study design, data collection, analysis, interpretation, writing, nor the decision to submit the article for publication.

We are grateful to two large consortia (SSGAC and CARDIoGRAMplusC4D) for publically sharing the genetic data we used in our causal analysis.

## REFERENCES

1. Hinkle LE, Jr., Whitney LH, Lehman EW, et al. Occupation, education, and coronary heart disease. Risk is influenced more by education and background than by occupational experiences, in the Bell System. Science 1968;16:238–46.

2. Manrique-Garcia E, Sidorchuk A, Hallqvist J, et al. Socioeconomic position and incidence of acute myocardial infarction: a meta-analysis. J Epidemiol Community Health 2011;65:301–9.

3. Veronesi G, Ferrario MM, Kuulasmaa K, et al. Educational class inequalities in the incidence of coronary heart disease in Europe. Heart 2016;102:958–65.

4. Davey Smith G, Hart C, Hole D, et al. Education and occupational social class: which is the more important indicator of mortality risk? J Epidemiol Community Health 1998;52:153–60.

5. Clark D, Royer H. The effect of education on adult mortality and health: evidence from Britain. Am Econ Rev 2013;103:2087–120.

6. Naess O, Hoff DA, Lawlor D, et al. Education and adult cause-specific mortality--examining the impact of family factors shared by 871 367 Norwegian siblings. Int J Epidemiol 2012;41:1683–91.

7. Davey Smith G, Ebrahim S. ‘Mendelian randomization’: can genetic epidemiology contribute to understanding environmental determinants of disease? Int J Epidemiol 2003;32:1–22.

8. Davey Smith G, Ebrahim S. What can mendelian randomisation tell us about modifiable behavioural and environmental exposures? BMJ 2005;330:1076–9.

9. Bowden J, Davey Smith G, Burgess S. Mendelian randomization with invalid instruments: effect estimation and bias detection through Egger regression. Int J Epidemiol 2015;44:512–25.

10. Bowden J, Davey Smith G, Haycock PC, et al. Consistent Estimation in Mendelian Randomization with Some Invalid Instruments Using a Weighted Median Estimator. Genet Epidemiol 2016;40:304–14.

11. Burgess S, Dudbridge F, Thompson SG. Re: “Multivariable Mendelian randomization: the use of pleiotropic genetic variants to estimate causal effects”. Am J Epidemiol 2015;181:290–91.

12. Okbay A, Beauchamp JP, Fontana MA, et al. Genome-wide association study identifies 74 loci associated with educational attainment. Nature 2016;533:539–42.

13. Deloukas P, Kanoni S, Willenborg C, et al. Large-scale association analysis identifies new risk loci for coronary artery disease. Nat Genet 2013;45:25–33.

14. Centers for Disease Control and Prevention (CDC). National Center for Health Statistics (NCHS). National Health and Nutrition Examination Survey Data. Hyattsville, MD: U.S. Department of Health and Human Services, Centers for Disease Control and Prevention, 1999-2014. http://www.cdc.gov/nchs/nhanes/, Accessed 2016, Aug 27.

15. Peasey A, Bobak M, Kubinova R, et al. Determinants of cardiovascular disease and other non-communicable diseases in Central and Eastern Europe: rationale and design of the HAPIEE study. BMC public health 2006;6:255. doi: 10.1186/1471-2458-6-255.

16. Evans A, Salomaa V, Kulathinal S, et al. MORGAM (an international pooling of cardiovascular cohorts). Int J Epidemiol 2005;34:21–7.

17. Zheng J, Erzurumluoglu AM, Elsworth BL, et al. LD Hub: a centralized database and web interface to perform LD score regression that maximizes the potential of summary level GWAS data for SNP heritability and genetic correlation analysis. Bioinformatics 2016. doi: 10.1093/bioinformatics/btw613. Epub ahead of print.

18. Bowden J, Del Greco MF, Minelli C, et al. Assessing the suitability of summary data for two-sample Mendelian randomization analyses using MR-Egger regression: the role of the 12 statistic. Int J Epidemiol 2016. doi: 10.1093/ije/dyw220. Epub ahead of print.

19. Baigi A, Holmen A, Hogstedt B, et al. Birthplace and social characteristics as risk factors for acute myocardial infarction in the province of Halland, Sweden. Public health 2002;116(5):279–84.

20. Lleras-Muney A. The relationship between education and adult mortality in the United States. Rev Econ Stud 2005;72:189–221.

21. Mazumder B. Does education improve health? A reexamniation of the evidence from compulsory schooling laws. Econ Perspect 2008;33:2–16.

22. Lager AC, Torssander J. Causal effect of education on mortality in a quasi-experiment on 1.2 million Swedes. Proc. Natl. Acad. Sci. USA 2012;109:8461–6.

23. Lundborg P. The health returns to schooling - what can we learn from twins? J Pop Econ 2013;26:673–701.

24. Madsen M, Andersen AM, Christensen K, et al. Does educational status impact adult mortality in Denmark? A twin approach. Am J Epidemiol 2010;172:225–34.

25. Arden R, Luciano M, Deary IJ, et al. The association between intelligence and lifespan is mostly genetic. Int J Epidemiol 2016;45:178–85.

26. Sondergaard G, Mortensen LH, Nybo Andersen AM, et al. Does shared family background influence the impact of educational differences on early mortality? Am J Epidemiol 2012;176:675–83.

27. Marioni RE, Ritchie SJ, Joshi PK, et al. Genetic variants linked to education predict longevity. Proc. Natl. Acad. Sci. USA 2016;113:13366–13371.

28. Stringhini S, Sabia S, Shipley M, et al. Association of socioeconomic position with health behaviors and mortality. JAMA 2010;303:1159–66.

29. Richmond RC, Hemani G, Tilling K, et al. Challenges and novel approaches for investigating molecular mediation. Hum Mol Genet 2016;25:R149–R56.

30. Varbo A, Benn M, Smith GD, et al. Remnant cholesterol, low-density lipoprotein cholesterol, and blood pressure as mediators from obesity to ischemic heart disease. Circ Res 2015;116:665–73.

31. Nelson CP, Hamby SE, Saleheen D, et al. Genetically determined height and coronary artery disease. N Engl J Med 2015;372:1608–18.

32. Fletcher JM. The promise and pitfalls of combining genetic and economic research. Health Econ 2011;20:889–92.

33. Rose G. Sick individuals and sick populations. Int J Epidemiol 1985;14:32–8.

34. Swerdlow DI, Preiss D, Kuchenbaecker KB, et al. HMG-coenzyme A reductase inhibition, type 2 diabetes, and bodyweight: evidence from genetic analysis and randomised trials. Lancet 2015;385:351–61.

